# Human monogamy in mammalian context

**DOI:** 10.1101/2025.08.19.671116

**Authors:** Mark Dyble

## Abstract

Monogamy has been argued to have played an important role in human evolution^1–5^ and, across animals more generally, evolutionary transitions to highly cooperative societies have been far more likely to occur in monogamous species^6–8^, raising the possibility that this may also have been the case for humans. However, the extent to which we can consider monogamy to be the species-typical human mating system is subject to debate^9–11^. Here, I provide comparative context on human mating behaviour by comparing the distribution of sibling types (full siblings versus half-siblings) across >100 human societies with equivalent data from 35 nonhuman mammal species. While cross-culturally variable, rates of full siblings in humans cluster closely with rates seen among socially monogamous mammals and fall consistently above the range seen in non-monogamous mammals. Although the human data is demonstrative of cross-cultural diversity in marriage and mating practices, the overall high frequency of full siblings is consistent with the characterisation of monogamy as the modal mating system for humans.

## Main Text

Evidence from birds^6^, mammals^8^, and insects^7,12^ all suggest that transitions to eusocial or cooperative breeding systems are more likely to occur in monogamous species, raising the possibility that the evolution of highly cooperative social behaviour in humans was also preceded by the evolution of monogamy^11,13^. Additionally, many derived features of human biology and behaviour have been hypothesised to have co-evolved with a monogamous mating system, including increased paternal care^14^, extended post-reproductive lifespans^1^, and the recognition of extensive networks of kin^2^.

Despite the hypothesised importance of monogamy in humans, the extent to which monogamy can be regarded our species-typical mating system has been subject to debate^9,11,13^. Although reduced sexual dimorphism^15^, reduced testes size^16^, and concealed ovulation^17^ in humans have all been argued to be indicative of a transition to reduced levels of polygyny during hominin evolutionary history^11^, our ability to reconstruct mating systems from the fossil record is limited and focus has instead been on making inferences from the ethnographic record.

Among contemporary or recent pre-industrial human societies, there is considerable diversity in marriage and mating norms and practices. For example, polygynous marriage (where a man is married to >1 woman at the same time) is permitted in ∼85% of a representative sample of pre-industrial societies^14,18^, leading to some to suggest that the predominance of monogamous marriage across much of the world today is an evolutionary novelty and a consequence of recent and rapid cultural change^9^. Others have argued that monogamy is a species-typical trait^19^ and point out that even within societies that permit polygynous marriage, the majority of marriages are still monogamous^18,20^. Further complexity is added to human mating and marriage practices by the occurrence of serial monogamy^21^, polyandrous marriages^22^ (where a woman is married to more than one man), partible paternity beliefs^23^, and extra-pair reproduction which is usually estimated to be <5%^24^ but can be much higher^25^.

An alternative but largely overlooked approach to assessing patterns of human mating is to consider the relative frequency full siblings, maternal half-siblings, and paternal half-siblings across populations^26^. At one extreme, exclusive lifetime reproductive monogamy (i.e. where an individual only ever reproduces with one other, who also only ever reproduces with them) would result in only full siblings, while random mating would result in many half-siblings and very few full siblings^27^.

Here, I analyse data from a global sample of 106 human populations (*n* = 197,695 sibling dyads in total) and compare this with a dataset of 35 nonhuman mammal species (*n* = 61,181 sibling dyads). Data on nonhuman mammal species was compiled from a systematic literature search building on cross-species analyses of reproductive skew^28^ and within-group relatedness^29,30^ (see Methods). The dataset includes 10 species categorised as socially monogamous (defined as a species where a reproductive pair share a common range and associate with each other for more than one breeding season^31^) and 25 species that are not socially monogamous (Supplementary Tables 1-2). The human data include ethnographic data on sibling proportions from ethnographic studies of kinship from 97 societies, including 80 from an existing dataset^26^ as well as data from ancient DNA (aDNA) analyses of the genetic relatedness among individuals excavated from nine archaeological sites ranging in age from the Avar-period in Central Europe ∼1,100 years ago^32^ to Neolithic Anatolia ∼8,000 years ago^33^ (Supplementary Table 1).

The proportion of siblings that are full siblings rather than paternal half-siblings or maternal half-siblings varied greatly across the human sample, from 26% among individuals excavated from an Early Neolithic site in Britain^34^ to 100% of siblings in four populations, including a Neolithic cemetery from Northern France^35^. Across the entire human sample, the average proportion of siblings that are full siblings (66%) was comparable to the rates seen in socially monogamous nonhuman mammals (mean = 70.6%, range = 45.2% to 100%, *n* = 10) and consistently exceeded the range seen in the non-monogamous nonhuman mammals (mean = 8.3% of siblings are full siblings, range 0-22%, *n* = 25, Figure 1). As such, human mating patters produce sibling distributions that very clearly cluster with socially monogamous species such as meerkats (*Suricata suricatta*, 59.0% full siblings) and African wild dogs (*Lycaon pictus*, 85.0% full siblings) rather than non-monogamous species such as chimpanzees (*Pan troglodytes*, 4.1% of siblings).

**Figure 1.**
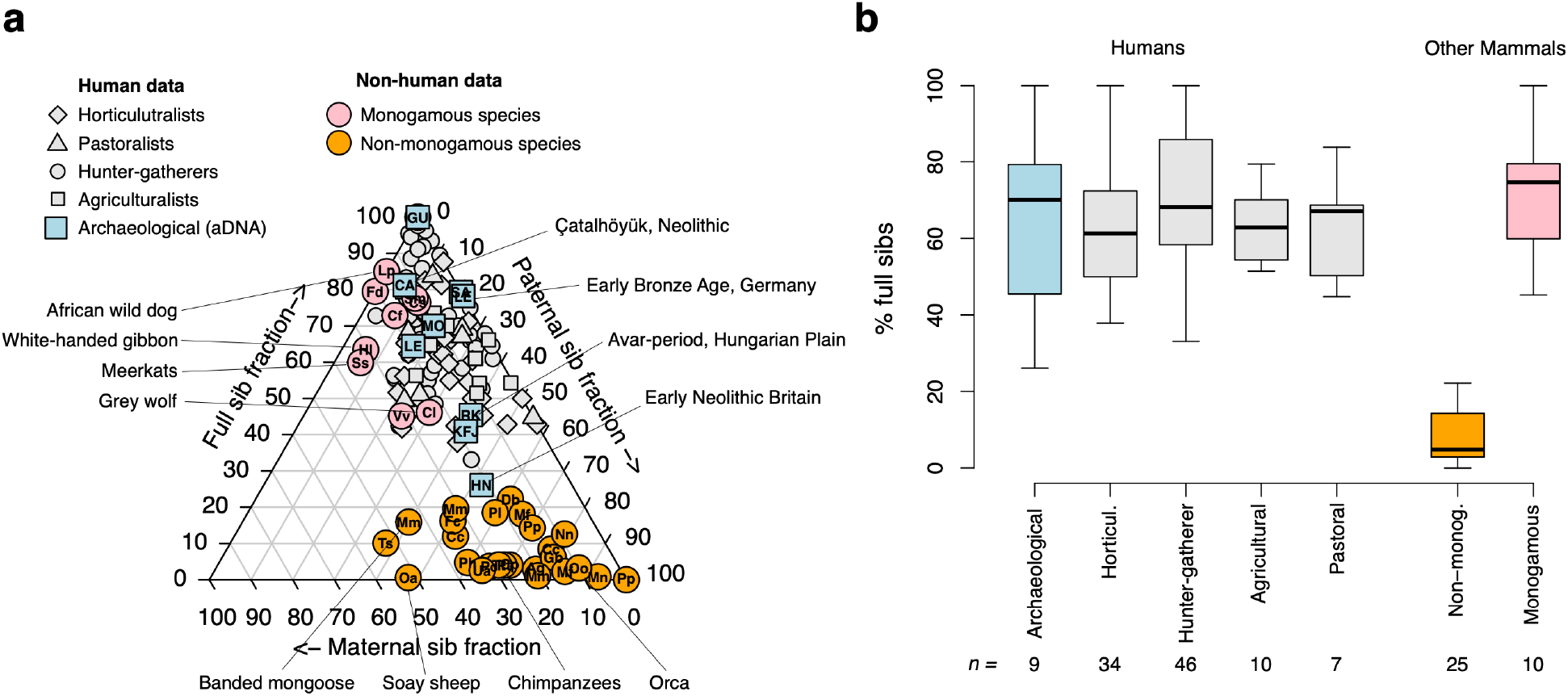
Human sibling proportions in mammalian context. **a**, ternary plot showing the proportion of full siblings, maternal half-siblings, and paternal half-siblings across the sample of human societies and nonhuman species. **b**, boxplot showing the proportion of full siblings across the sample of human and nonhuman species. Boxplots show median values, 50th percentile values (box outline) and range (whiskers). Colours correspond to ancient human data (blue), ethnographic human data (grey), non-monogamous nonhuman mammals (orange), and monogamous nonhuman mammals (pink). Letters in circles are abbreviated species names (e.g. Oa = *Ovis aries*, Soay sheep, see Supplementary Tables 3-4).

Although it is clear that random mating will result in very few full siblings and that exclusively monogamous mating will result in only full siblings, the nature of the relationship between extra-pair paternity and the proportion of full siblings is not obvious. In order to better understand this relationship, I constructed a computational model in which I varied rates of mating within a simulated population between exclusive monogamy (extra-pair paternity (*e*) = 0) and random mating (*e* = 1) (Methods). Varying rates of *e* in this scenario produces a non-linear negative relationship between *e* (characterised as ‘extra-pair paternity’) and the proportion of full siblings (Figure 2a), with relatively modest rates of extra-pair paternity having a disproportionate effect on the production of half-siblings. For example, a 25% extra-pair paternity rate will result in only of ∼40% sibling dyads being full siblings, while 50% extra-pair paternity will result in ∼15% of siblings being full siblings (Fig 2a).

**Figure 2.**
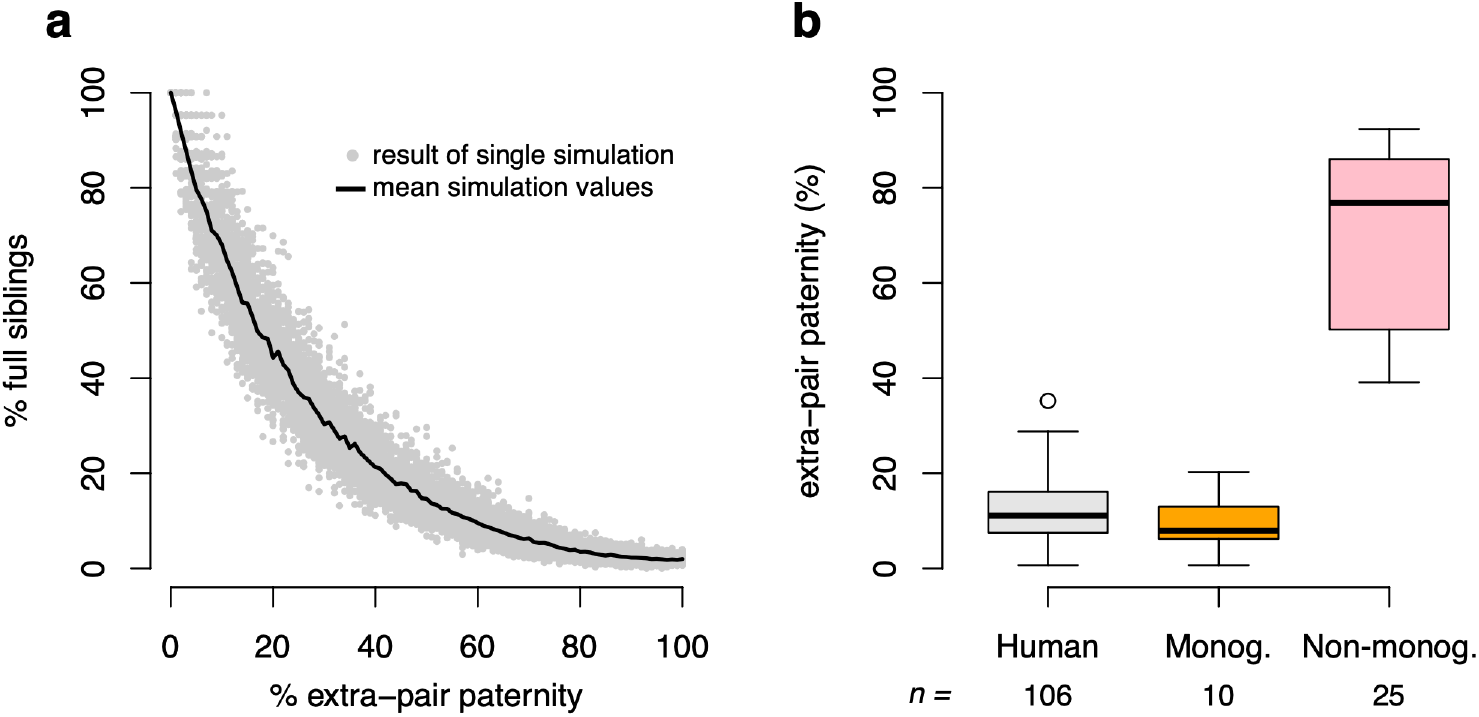
Estimating extra-pair paternity from proportions of full siblings. **a** simulation results to estimate the proportion of siblings that are full siblings given varying rates of extra-pair paternity. Note that when *e* is high, the description of the parameter as extra-pair paternity is stretched because stable reproductive pairs cease to be present. Nonetheless, *e* captures the spectrum between exclusive monogamy and random mating. **b** estimated *e* for human and nonhuman mammal samples based on extrapolating observed sibling proportions through the predictions of the theoretical model.

Although the exact relationship between extra-pair paternity and full sibling production is also influenced by group size, parity, and extra-group mating, the relationship is largely robust to these differences (see Extended Data Figure 2). Extrapolating back from the oberved rates of full siblings (Figure 2b), we can estimate *e* (extra-pair paternity) across the human sample to vary from 0 to 35%, averaging 12% (Figure 2c). Mean estimated rates produced for the monogamous and non-monogamous species are 9.9% and 69%, respectively (Figure 2b).

A final means of comparing reproductive behaviour in humans with that of other species is to estimate rates of reproductive monogamy directly. This is here defined and measured as the proportion of reproductive individuals who have only reproduced with one individual and where that individual has themselves only reproduced with the focal individual. This is possible for a sub-sample of the total dataset for which parentage data are available (see Methods). An average of 61.0% of individuals across the ethnographic human datasets were reproductively monogamous (*n* = 17 societies). Similarly, an average of 69.1% individuals across the sufficiently large archaeological samples (*n* = 4) were reproductively monogamous. These rates are higher than seen in the nonhuman monogamous species (38.6%, *n* = 5) and much higher than seen in non-monogamous species (9.2%, *n* = 14, Figure 2).

## Discussion

It has been hypothesised elsewhere that monogamy has played an important role in human evolution^2,3,13,36^, as well as in the evolution of highly social animal societies more broadly^6–8,12^. While previous work has relied on inferences from the fossil record^15^, or cross-cultural comparisons of marriage norms^10,18^, I here focus on the measuring the outcome of mating systems: the relative proportion of full and half-siblings born into a population; a direct, theoretically salient but relatively overlooked approach^26^. Although the variation in sibling composition across the human dataset is demonstrative of the obvious fact that humans are not *universally* monogamous, the finding that human rates of full siblings overlap with the range seen in socially monogamous mammal species and are consistently higher than those observed in non-monogamously mating mammal species lends weight to the broad-stroke characterisation of monogamy as the modal mating pattern for our species.

If we can regard humans as a typically monogamous species, then we join the estimated minority of ∼9% of mammals that are socially monogamous^31^. However, where humans deviate from the vast majority of socially monogamous mammal species is that we live in social groups in which multiple females breed: socially monogamous mammalian species tend to either live in groups containing only the breeding pair and their offspring or in singular breeding groups where only one female of many reproduces^8^. Although monogamous pairs sometimes aggregate in a few species, the only other species of mammal that has been suggested live in stable multi-male multi-female groups containing multiple monogamous pairs is the Patagonian mara (*Dolichotis patagonum*), where large numbers of monogamous pairs may share a warren^31,37^. Humans also differ from most socially monogamous species in being monotocous^38^ (i.e. typically producing single offspring per pregnancy rather than litters); all of the monogamous mammal species in the present sample, with the exception of the white-handed gibbon, are litter-bearing (polytocous) species (Supplementary Table 4). Therefore, while humans are not unusual among mammals in producing large numbers of full siblings through monogamous mating, we are unusual in doing so while being monotocous and living in plural breeding multi-male multi-female groups. Given that all other African great apes live in groups and have either have polygynous or polygynandrous mating systems, it is probable that human monogamy evolved from a non-monogamous group-living state, a transition that is highly unusual among mammals more generally^31^ and which suggests different pressures selected for the evolution of monogamy in humans as compared to other species, possibly related to the energetic demands of our large brains and slow growth^39,40^.

The “monogamy hypothesis” holds that the evolution of highly cooperative social behaviour is more likely to occur in species in which parents mate monogamously and produce full siblings^12^. The relevance of full siblings is that, in diploid species, individuals can expect to be as genetically related to their full siblings as they would be to their own offspring (*r* = 0.5), more readily favouring altruistic behaviour toward kin and helping parents to raise younger siblings (‘helping at the nest’). The theoretical predictions about the relationship between extra-pair paternity and full sibling production shown here offer a refinement to this hypothesis because the non-linear relationship between extra-pair paternity and the production of full siblings suggests that even modest deviations from monogamy can have a large influence on diluting the proportion of siblings that are full siblings (e.g. ∼25% extra-pair paternity leads to ∼40% siblings as full siblings, though noting that the overall number of siblings of any kind also increases). This could go some way to explaining why, for example, the proportion of birds that are cooperative breeders (∼9%) is so much lower than the proportion that are socially monogamous (∼90%)^41^: it is not uncommon for socially monogamous birds have estimated rates of extra-pair paternity exceeding 20%^42^; this may sufficiently reduce relatedness to make transitions to cooperative breeding unlikely.

In humans, the majority of the half-siblings are paternal half-siblings rather than maternal half-siblings^26^. This is indicative of higher levels of male reproductive skew than female reproductive skew in humans^26,28^ and is consistent with polygynous marriages being much more commonly permitted across human societies than polyandrous marriages, which are rare^18,43^. Since the ethnographic data are collected through self-reported parentage (see Methods) rather than genetic analysis, it is possible that there is some misattributed paternity within the ethnographic human dataset. However, the general consistency between genetic and ethnographic datasets in the frequency of full siblings suggests the effect may not be great and where extra-pair paternity has been estimated in human societies it is typically well under 5%^24^. Although a recent finding has reported 46% extra-pair paternity among Himba pastoralists^25^, this is a considerable outlier and was also accompanied by high paternity confidence; men were able to correctly identify whether they were the biological father of their spouse’s children in ∼75% of cases.

It is important to note that what has been measured by analysing sibling proportions is patterns of reproduction, rather than patterns of mating and is therefore an assessment of reproductive monogamy, rather than monogamy in mating. In most mammals, mating and reproductive patterns will usually being tightly linked. In humans, this link will be diminished in part by birth control; not just the highly effective contraceptives developed over the last century but also ‘natural’ birth control methods and cultural practices to control fertility that are a consistent feature cross-culturally^44,45^.

Human marriage and mating practices are highly variable, and there has been much interest in explaining this variability from an evolutionary perpective^9,46,47^. Here, I focus not on the variation but on general features of human mating in comparative mammalian context, concluding that monogamy can be regarded as the species-typical system for *Homo sapiens*. The value of such a generalisation is that it helps contribute to the assessment of the plausibility of hypotheses that have situated monogamy as a core human characteristic that has co-evolved with various aspects of our biology and behaivour^1,2,5,39^. If these hypotheses are correct, monogamy has played an important role in human evolution, allowing increased energetic investment and extended juvenility through paternal care that allow us to have more offspring at shorter intervals^39^ and to establish the extended kinship networks (through the identification of paternal and affinal kinship^19,48^) that provided the first step in building large-scale societies and cultural networks that have been crucial to our success as a species^49^.

## Methods

### Human dataset

The human sample includes sibling proportions data from 106 human populations derived through archaeological or ethnographic data. The archaeological data are based on ancient DNA (aDNA) analyses of kinship from archaeological sites (Supplementary Table 1)^32–35,50–52^. To assemble this dataset, I conducted a literature search of journal articles on Web of Science with the search terms ‘(kinship OR relatedness OR genealogy) AND (aDNA OR ancient OR archaeological)’. I considered all papers returned up to and including July 2025 that included an arbitrary sample size threshold of at least 15 sibling dyads. This resulted in a sample of seven publications, two of which contained data on two separate archaeological sites with >15 sibling dyads: Wang et al.^32^ included data from Avar-period cemeteries at Leobersdorf (90 sibling dyads) and Mödling-An der Goldenen Stiege (294 sibling dyads) while Gnecchi-Ruscone et al.^51^ contained data from Rákóczifalva cemetery (165 sibling dyads) and Kunszallas-Fulopjakab cemetery (71 sibling dyads). Sibling relationships were calculated through assembling parentage lists from genealogical diagrams provided in the publications (Supplementary Table 1). Only sibling relationships between sampled (rather than genealogically inferred) individuals were included.

The ethnographic data are built on genealogies compiled by ethnographers from a global sample of 97 pre-industrial human societies engaged in a range of subsistence types. Data on numbers of siblings for 80 of the 97 societies were included in an analysis by Ellsworth et al.^26^ who compiled data through the open access *kinsources* database (kinsources.net) (N = 56 of 80), with additional information from two other publications^4,53^. I use the sibling proportion data tabulated by Ellsworth et al.^26^ as well as sibling numbers for an additional 17 societies based on analysis of kinship datasets that have been made available through the *kinsources* project since 2016 (Supplementary Table 2). The categorisation of subsistence types was based on literature review and cross-referencing the society names with the DPLACE ethnographic database^54^. Although categorical distinctions between subsistence types are often somewhat arbitrary^55^, such categories are used here only to give an indication of economic diversity within the sample, rather than to test hypotheses about the relationship between dominant subsistence mode and sibling proportions.

The similar proportions between the human genetic (*n* = 9, mean full siblings = 65%) and genealogical data (*n* = 97, mean full siblings = 66%) do not suggest a major discrepancy between the genealogically derived ethnographic and genetically derived archaeological datasets. Nonetheless, it is likely that some of the reported paternal relationships in the ethnographic dataset do not match genetic relatedness. To estimate the effect of possible underestimated extra-pair paternity on the results, I took the sample of 17 societies from the additional ethnographic dataset included here (Supplementary Table 2) and simulated extra-pair paternity by randomly replacing father names with those of a randomly sampled father at a given rate. The baseline proportion of full siblings averaging across these 17 societies in 69%, close to the overall mean for the ethnographic dataset of 66%. A simulated 1% extra-pair paternity rate reduces the proportion of siblings that are full siblings to 66.1%, a 3% extra-pair paternity rate reduces this to 62.6% and a 10% extra-pair paternity rate reduces this to 51.8% (Extended Data Figure 1). In all these scenarios these rates remain significantly higher than the rates seen in the non-monogamous mammal sample (range 0-22% full siblings, Extended Data Figure 1).

### Nonhuman mammal dataset

While the mating systems of many of the >∼6,000 mammals species have been qualitatively classified^31^, data on sibling proportions requires genetic parentage analysis to have been conducted, something which has been done for a much more modest number of species. To restrict the sample to mammal species for which genetic data have been collected and where sibling data might therefore be available, I compiled a list of species that had been included in recent comparative studies of mammalian reproductive skew^28^ and kinship composition^29,30^ based on genetic data (*n* = 70 species in total). For each of these candidate species, I followed up the references provided in the prior studies and complemented this with a Web of Science search for ‘[relevant species binomial] AND (siblings OR kinship OR paternity)’. These searches yielded 35 species for which sibling numbers could be determined, either because they were directly reported or because a parentage list was provided from which sibling distributions could be estimated (Supplementary Tables 3-4). In all cases, data come from a single study population with the exception of chimpanzees (*Pan troglodytes*) for which data were available for four populations (Supplementary Table 3; the species-level sibling numbers for chimpanzees are the sum of the numbers across these four populations). For categorical data on whether a species was socially monogamous, a plural or singular breeder, and monotocous or polytocous, species names were cross-referenced with published comparative analyses of plural breeding and monogamy in mammals^31,38^.

### Estimating sibling proportions and reproductive exclusivity

Sibling proportions were estimated by calculating the numbers of individuals of known parentage who shared two parents (full siblings), a mother but had different fathers (maternal half-siblings), or a father but different mothers (paternal half-siblings). Reproductive exclusivity was calculated as the proportion of reproducing individuals (i.e., those who were represented in the dataset as the mother or father of at least one individual) who had reproduced with only one other individual who had themselves only reproduced with the focal individual. For this analysis the sample was necessarily restricted to those populations or species for which parent lists were available (all archaeological datasets, 17/97 ethnographic datasets, 20 of 35 nonhuman species datasets) with the five archaeological sites featuring fewer than 10 parents also excluded.

### Modelling sibling proportions

The computational model was written in R^56^ using custom code. The model starts by considering a group containing *N_f_*females who produce *k* offspring and then varying the extent to which they produce these offspring with one male versus multiple males. In the purely monogamous scenario (extra-pair paternity (*e*) = 0) there is exclusively monogamous mating and all siblings are full siblings. In the totally promiscuous scenario (*e* = 1) males and females are randomly paired to reproduce and full siblings are rare. At intermediate rates (0 > *e* < 1), we start as in the exclusively monogamous condition and the father identity of each individual is then sampled to a random male with probability *e*. The model assumes a balanced sex ratio of potential mothers and fathers as well as all paternities falling within the group (though for relaxation of this assumption see Extended Data Figure 2). Estimation of *e* in the empirical data is based on taking the proportion of full siblings observed in each population/species and finding the simulation results that produced a proportion of full siblings within 2 percentage points of the observed proportion and taking the average value of *e* for those simulations.

## Data availability

Sibling proportions for human populations and nonhuman species are provided in Supplementary Tables 1-4 and also available in an electronic format on GitHub. Where data on sibling proportions are constructed from parent lists, these are also available on GitHub (https://github.com/mdyble/siblings.git).

## Code availability

Code for the estimation of sibling proportions based on parentage lists, production of figures, and simulations of sibling proportions are available on GitHub (https://github.com/mdyble/siblings.git).

## Author contributions

M.D. compiled and analysed the data and model and wrote the paper.

