## Supplementary Materials for "Human monogamy in mammalian context"

**Supplementary Material for 'Human monogamy in mammalian context'**

Author: Mark Dyble

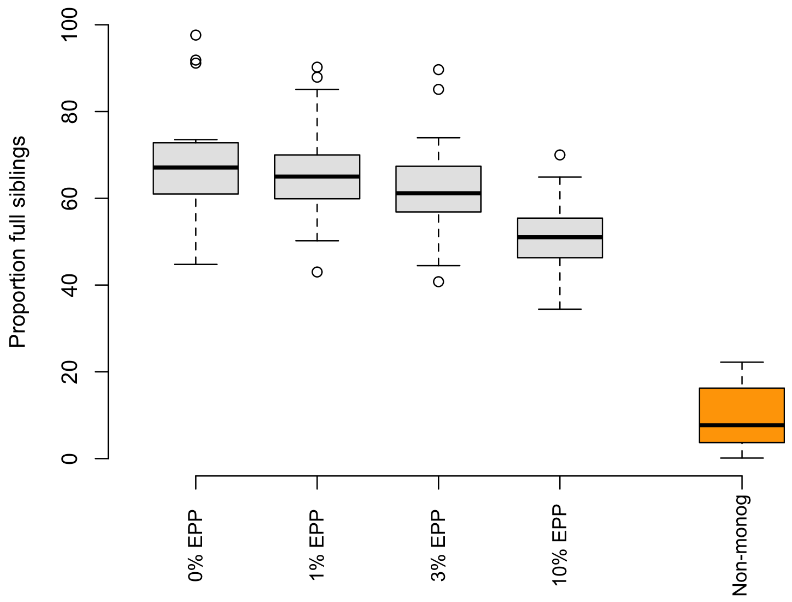


**Extended Data Figure 1 | Proportion of full siblings estimated across a sample of 17 populations given simulated levels of extra-pair paternity.** Even with a high degree of extra-pair paternity, rates of full siblings remain consistently higher than observed in the non-monogamous mammal sample.

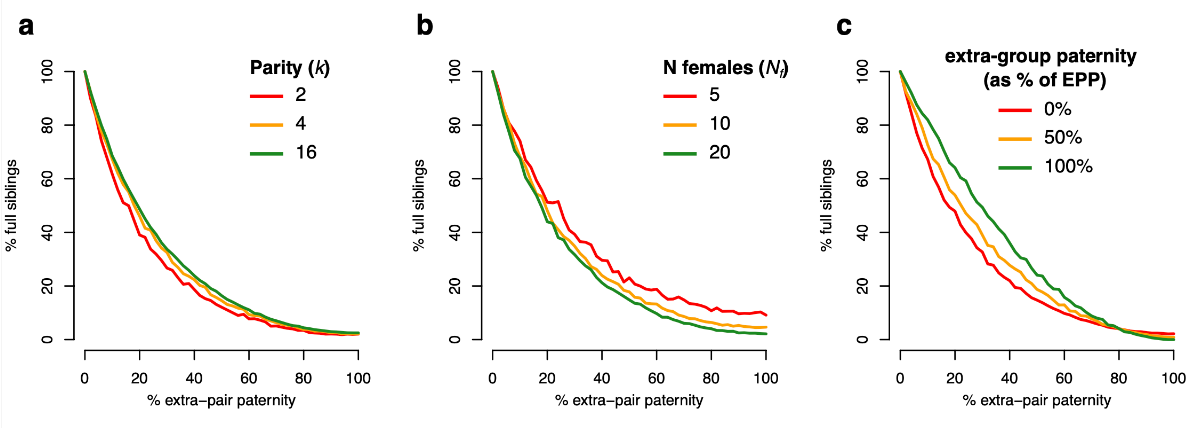


**Extended Data Figure 2 | Robustness checks for simulations estimating the proportion of full siblings produced across varying degrees of extra pair-paternity. a.** effect of varying parity (i.e. the number of offspring born to each female) on rates of full siblings. **b**. effect of number of females on rates of full siblings. **c.** effect of extra-group paternity on full sibling rates. This is implemented by replacing an extra-pair father with a random m1ale extra-pair father from outside the group at a given rate (here, 0%, 50%, 100%). Unless otherwise stated, $Nf = 20$ and $k = 4$.

**Supplementary Table 1: Sibling numbers from archaeological sites.**

| Description | Full siblings | Paternal half sibs | Maternal half sibs | Reference |
|---|---|---|---|---|
| Gurgy ‘les Noisats’ burial site. European Neolithic, ~3,000–6,500-y-old. | 44 | 0 | 0 | Rivollat et al. 2023[1] From Figure 2. Text confirms absence of half-sibling dyads. |
| Leubingen burial ground. Early Bronze Age, Germany. ~4,000-y-old | 18 | 5 | 0 | Penske et al. 2024[2] From Figure 2 |
| Rákóczifalva cemetery, Avar-period, Hungarian plain. ~570 to the mid-ninth century. | 75 | 66 | 24 | Gnecchi-Ruscone et al. 2024[3] From Supplementary Fig. 10 |
| Kunszallas-Fulopjakab cemetery, Avar-period, Hungarian plain. Mid-seventh-century AD | 35 | 25 | 11 | Gnecchi-Ruscone et al. 2024[3] From Extended Data Fig. 1 |
| Bronze Age Srubnaya-Alakul cultural tradition, Southern Urals. ~3,800-y-old burial mound | 23 | 6 | 0 | Blöcher et al. 2023[4] From Figure 2 |
| Hazleton North long cairn, Early Neolithic Britain. ~5,700-y-old | 6 | 12 | 5 | Fowler et al. 2022[5] From Figure 1 |
| Çatalhöyük East Mound, Neolithic Anatolia, ~8000 to 9000-y-old | 13 | 1 | 2 | Yüncü et al. 2025[6] From Figures S15-S20 |
| Leobersdorf cemetery Avar-period. Seventh to early ninth century CE. | 58 | 15 | 17 | Wang et al. 2025[7] Figure 2 |
| Mödling-An der Goldenen Stiege cemetery, Avar-period. Seventh to early ninth century CE. | 206 | 55 | 33 | Wang et al. 2025[7] Figure 3 |

**Supplementary Table 2: Sibling numbers from the additional ethnographic sources available on the *kinsources* publicly available dataset** (https://www.kinsources.net)**.** Predominant subsistence categories defined by cross-reference with DPLACE ethnographic database or on literature review. Society names follow those used on *kinsources*.

| **Society [Region]** | **Predominant subsistence** | **Full sibs** | **Paternal half sibs** | **Maternal half sibs** |
|---|---|---:|---:|---:|
| Baruya [New Guinea] | Horticultural | 7011 | 3810 | 773 |
| Bwa [West Africa] | Agricultural | 29443 | 16067 | 2769 |
| Dogon [West Africa] | Agricultural | 22016 | 16414 | 4358 |
| Duu Rea [North/Central Africa] | Pastoral | 3650 | 1374 | 252 |
| Jie [East Africa] | Agricultural | 86 | 44 | 0 |
| Kel Owey [West Africa] | Pastoral | 11180 | 4510 | 974 |
| Kodiak [North America] | Hunter-gatherer | 806 | 39 | 262 |
| Mebengokre (Kayapo) [Brazil] | Agricultural | 1846 | 430 | 235 |
| Mowanjum [Australia] | Hunter-gatherer | 82 | 1 | 1 |
| Murriny-Patha [Australia] | Hunter-gatherer | 46 | 23 | 3 |
| Nucoorilma (Tingha) [Australia] | Hunter-gatherer | 1613 | 94 | 49 |
| Nunivak [North America] | Hunter-gatherer | 806 | 211 | 199 |
| Samburu [East Africa] | Pastoral | 607 | 749 | 0 |
| Sarmi [New Guinea] | Hunter-gatherer | 229 | 170 | 36 |
| Tiwi [Australia] | Hunter-gatherer | 7 | 2 | 1 |
| Tlingit [North America] | Hunter-gatherer | 103 | 5 | 5 |
| Todas [India] | Pastoral | 1621 | 324 | 433 |

**Supplementary Table 3: Species included in study, with references and notes.** Prior study lists the previous comparative studies in which this species was included; R = Ross et al. 2023[8], P = Pereira et al. 2023[9], D = Dyble & Clutton-Brock 2020[10]. Search for information on sibling numbers were made for all species listed here. Those indicated as not included in the present study are those for which sufficient information could not be found.

| Species | Common name | Prior study | Present study |
|---|---|---|---|
| *Antechinus agilis* | Agile antechinus | R | No |
| *Antechinus stuartii* | Brown antechinus | R | No |
| *Arctocephalus gazella* | Antarctic fur seal | R | Yes, Table S1 from Bonin et al. 2014[11] |
| *Artibeus jamaicensis* | Jamaican fruit bat | P | No |
| *Brachyteles hypoxanthus* | Northern muriqui | R | No |
| *Bradypus variegatus* | Brown-throated sloth | R | No |
| *Canis lupus* | Grey wolf | P | Yes, Figure 1 Hedrick et al. 2014[12] |
| *Canis rufus* | Red wolf | R | No |
| *Canis simensis* | Ethiopian wolf | D | Yes, in text of Randall et al. 2007[13] but only full versus half |
| *Capreolus capreolus* | Roe deer | R | No |
| *Castor fiber* | Eurasian beaver | R | Yes, Table 2 of Nimje et al. 2019[14] |
| *Cebus capucinus* | White-faced capuchin | R,P | Yes, text of results in Godoy et al. 2016[15] |
| *Ceratotherium simum* | White rhinoceros | R | No |
| *Cercopithecus mitis* | Blue monkey | P | No |
| *Cervus elaphus* | Red deer | R | No |
| *Chlorocebus pygerythrus* | Vervet monkey | R | Yes, Table S3 Minkner et al. 2018[16] |
| *Crocuta crocuta* | Spotted hyena | R,P,D | Yes, Numbers given in Figure 3 of Wahaj et al. 2004[17] |
| *Ctenodactylus gundi* | Common gundi | P | No |
| *Cuon alpinus* | Dhole | P | No |
| *Cynomys ludovicianus* | Black-tailed prairie dog | P | No |
| *Cynopterus sphinx* | Indian fruit bat | R | No |
| *Diceros bicornis* | Black rhinoceros | R | Yes, Table 2 of Garnier et al. 2001[18] |
| *Eptesicus fuscus* | Big brown bat | P | No |
| *Equus caballus* | Wild horse | R | No |

| | | | |
|---|---|---|---|
| *Equus quagga* | Plains zebra | R | No |
| *Felix catus* | Feral cat | R | Yes, Based on parentage data given in Table 3 of Natoli et al. 2007[19] |
| *Fukomys damarensis* | Damaraland mole rat | R,P,D | Yes, Table 2 of Burland et al. 2002[20] |
| *Gorilla beringei* | Mountain gorilla | R | Yes, sibling numbers given in results of Grebe et al. 2002[21] |
| *Hylobates lar* | White-handed (or lar) gibbon | R | Yes, Table IV of Barelli et al. 2013[22] |
| *Leontopithecus rosalia* | Golden lion tamarin | R | No |
| *Lepus americanus* | Snowshoe hare | R | No |
| *Loxodonta africana* | African elephant | R | No |
| *Lycaon pictus* | African wild dog | R,D | Yes, Figure 3 in Girman et al. 1997[23] lists 68 full sibs pairs and 12 half-sib pairs. Main text clarifies that all half sibs shared a mother but not a father. |
| *Macaca fascicularis* | Long-tailed macaque | P | Yes, Figure 2 of Ruiter & Geffen 1998[24] |
| *Macaca fuscata* | Japanese macaque | R | Yes, Table S4 of Ishizuka et al. 2024[25] |
| *Macaca mulatta* | Rhesus macaque | R,P | Yes, given on p39 as a per individual rate in Widdig 2003[26] |
| *Macaca nigra* | Crested macaque | R | Yes, ESM of Engelhardt et al. 2016[27] |
| *Macaca sylvanus* | Barbary macaque | R | No: Brauch et al. 2008[28] include data but only 12 individuals and 14 sibships |
| *Marmota flaviventris* | Yellow-bellied marmot | P | No |
| *Meles meles* | European badger | R | Yes, Figure 2 of Dugdale et al. 2008[29] |
| *Mirounga angustirostris* | Elephant seal | R | No |
| *Mungos mungo* | Banded mongoose | D | Yes, Table S1.2.7 in Sanderson et al. 2015[30] |
| *Myocastor coypus* | Coypu | D | No |
| *Myotis bechsteinii* | Bechstein's bat | R,P | No |
| *Nasua nasua* | Ringtailed coati | R | Yes, Based on parentage data in Table 2 of Hirsch & Maldonado 2010[31] |
| *Odocoileus virginianus* | White-tailed deer | R | No |
| *Orcinus orca* | Orca | R,P | Yes, Based on Table S6 Ford et al. 2018[32] |

| | | | |
|---|---|---|---|
| *Ovis aries* | Soay sheep | R | Yes, listed in Table 2 Morrissey and Wilson 2010[33] |
| *Ovis canadensis* | Bighorn sheep | R | No |
| *Pan paniscus* | Bonobo | R | Yes, Tables 2 and 3 in Ishizuka et al. 2018[34] |
| *Pan troglodytes* | Common chimpanzee | R,P,D | Yes, Mahale: Table 1 in Inoue et al. 2008[35] (0Full, 7PH, 0MS)<br><br>Tai: Fig 2 in Vigilant et al 2001[36] (0 full, 55PH, 16MH)<br><br>Budongo: Table 1, Newton-Fisher et al. 2010[37] (2Full, 64PH, 16MH)<br><br>Gombe: Table A1 from Wroblewski et al. 2009[38] (12Full, 110PH, 62MH) |
| *Panthera leo* | African lion | R,D | Yes, Parentage data from Figure 5 from Gilbert et al. 1991[39] |
| *Papio cynocephalus* | Savannah baboon | R,P | Yes, Table A1 Alberts et al. 2006[40] |
| *Papio hamadryas anubis* | Olive baboons | R | Yes, text of Lynch et al. 2017[41] |
| *Pecari tajacu* | Collared peccary | R | No |
| *Peromyscus californicus* | California deermouse | R | Yes, Figure 1 from Ribble 1991[42] |
| *Petrogale penicillata* | Rock wallaby | R | Yes, from Appendix of Hazlitt et al. 2006[43] |
| *Phascolarctos cinereus* | Koala | R | No |
| *Phoca vitulina* | Harbour seal | R | No |
| *Propithecus verreauxi* | Sifaka | R,D | No |
| *Rupicapra rupicapra* | Northern chamois | R | No |
| *Saguinus mystax* | Moustached tamarin | D | Yes, Table 3 Huck et al. 2005[44] |
| *Stenella frontalis* | Atlantic spotted dolphin | R | No |
| *Suricata suricatta* | Meerkat | R,D | Yes, personal communication with C. Duncan |
| *Tamias amoenus* | Yellow-pine chipmunk | R | No |
| *Tamias striatus* | Eastern chipmunk | R | Yes, Online data for Frappier-Lecomte et al. 2025[45] |

| | | | |
|---|---|---|---|
| *Tursiops truncatus* | Bottlenose dolphin | R | Yes, Table S2 in Wiszniewski et al. 2012[46] |
| *Ursus americanus* | Black bear | R | Yes, Table 2 in Kovach and Powell 2003[47] |
| *Vulpes vulpes* | Red fox | P | Yes, Table 1 from Iossa et al. 2009[48] |
| *Zalophus wollebaeki* | Galapagos sea lion | R | No |

**Supplementary Table 4**: Sibling proportions across mammals. Plural/singular breeding and monotocous/polytocous categories based on data from Lukas and Clutton-Brock (2020)[49]. Social monogamy category cross-referenced with dataset from Lukas and Clutton-Brock (2013)[50]. [1]Species absent from Lukas and Clutton-Brock dataset, own coding decision made. [2]Categories used are those listed for *Petrogale assimilis* which is part of the *P. lateralis/penicillata* species complex. [3]Listed as *Cercopithecus aethiops.* [4]Listed as *Cryptomys damarensis*. [5]Not listed but classified as non-monogamous, like all *Felix* species. [6]78 full siblings and 24 half-siblings listed in reference but not broken down by maternal/paternal half-siblings; assumption made of an equal split of maternal and paternal half siblings.

| Species | Full sibs | Paternal half sibs | Maternal half sibs | % Full | Breeding | Social Monogamy |
|---|---|---|---|---|---|---|
| *Arctocephalus gazella* (Antarctic fur seal) | 9 | 238 | 63 | 2.9 | Plural, monotocous | No |
| *Canis lupus* (Grey wolf) | 542 | 348 | 284 | 46.2 | Singular, polytocous | Yes |
| *Canis simensis* (Ethiopian wolf) | 78 | 12[6] | 12[6] | 76.5 | Singular, polytocous | Yes |
| *Castor fiber* (Eurasian beaver) | 371 | 41 | 97 | 72.9 | Singular, polytocous | Yes |
| *Cebus capucinus* (White-faced capuchin) | 75 | 689 | 123 | 8.5 | Plural, monotocous | No |
| *Chlorocebus pygerythrus* (Vervet monkey) | 10 | 173 | 64 | 4 | Plural, monotocous | No[3] |
| *Crocuta Crocuta* (Spotted hyena) | 12 | 53 | 35 | 12 | Plural, polytocous | No |
| *Diceros bicornis* (Black rhinoceros) | 12 | 33 | 9 | 22.2 | Singular, monotocous | No |
| *Felix catus* (Feral cat) | 32 | 99 | 66 | 16.2 | Singular, polytocous[1] | No[5] |
| *Fukomys damarensis* (Damaraland mole rat) | 3154 | 0 | 813 | 79.5 | Singular, polytocous | Yes[4] |
| *Gorilla beringei* (Mountain gorilla) | 43 | 555 | 101 | 6.2 | Plural, monotocous | No |
| *Hylobates lar* (White-handed gibbon) | 33 | 3 | 16 | 63.5 | Singular, monotocous | Yes |
| *Lycaon pictus* (African wild dog) | 68 | 0 | 12 | 85 | Singular, polytocous | Yes |

| | | | | | | |
|---|---|---|---|---|---|---|
| *Macaca fascicularis* (Long-tailed macaque) | 17 | 62 | 15 | 18.1 | Plural, monotocous | No |
| *Macaca fuscata* (Japanese macaque) | 2 | 74 | 12 | 2.3 | Plural, monotocous | No |
| *Macaca mulatta* (Rhesus Macaque) | 0.138(%) | 9.887(%) | 2.62(%) | 1.1 | Plural, monotocous | No |
| *Macaca nigra* (Celebes crested macaque) | 2 | 219 | 15 | 0.8 | Plural, monotocous | No |
| *Meles meles* (European badger) | 147 | 370 | 234 | 19.6 | Plural, polytocous | No |
| *Mungos mungo* (Banded mongoose) | 3595 | 9003 | 9994 | 15.9 | Plural, polytocous | No |
| *Nasua nasua* (Ring-tailed coati) | 105 | 656 | 71 | 12.6 | Plural, polytocous | No |
| *Orcinus orca* (Killer whale) | 15 | 400 | 45 | 3.3 | Plural, monotocous | No |
| *Ovis aries* (Soay sheep) | 146 | 11496 | 12611 | 0.6 | Plural, monotocous[1] | No |
| *Pan paniscus* (Bonobo) | 0 | 18 | 0 | 0 | Plural, monotocous | No |
| *Pan troglodytes* (Chimpanzee) | 14 | 236 | 94 | 4.1 | Plural, monotocous | No |
| *Panthera leo* (African lion) | 15 | 48 | 18 | 18.5 | Plural, polytocous | No |
| *Papio cynocephalus* (Savannah baboon) | 31 | 551 | 262 | 3.7 | Plural, monotocous | No |
| *Papio hamadryas anubis* (Olive baboon) | 4 | 50 | 30 | 4.8 | Plural, monotocous | No |
| *Peromyscus californicus* (California deermouse) | 82 | 0 | 0 | 100 | Singular, polytocous | Yes |
| *Petrogale penicillate* (Rock wallaby) | 37 | 182 | 40 | 14.3 | Singular, monotocous[2] | No[2] |
| *Saguinus mystax* (Moustached tamarin) | 59 | 8 | 9 | 77.6 | Singular, polytocous | Yes |
| *Suricata suricatta* (Meerkat) | 59.9(%) | 6.3(%) | 33.8(%) | 59.9 | Singular, polytocous | Yes |
| *Tamias striatus* (Eastern chipmunk) | 136 | 537 | 743 | 9.6 | Singular, polytocous | No |
| *Tursiops truncates* (Bottlenose dolphin) | 2 | 33 | 14 | 4.1 | Plural, monotocous | No |
| *Ursus americanus* (Black bear) | 2 | 50 | 26 | 2.6 | Singular, polytocous | No |
| *Vulpes vulpes* (Red fox) | 75 | 39 | 52 | 45.2 | Singular, polytocous | Yes |

**Supplementary Material References**

26. Widdig, A. Paternal kinship among adult female rhesus macaques (Macaca mulatta). (Humboldt-Universität zu Berlin, 2003).
